# Reactivity to conditioned threat cues is distinct from exploratory drive in the elevated plus-maze

**DOI:** 10.1101/2022.03.21.485161

**Authors:** Joe R Hilton, Susannah R Simpson, Emily R Sherman, Will Raby-Smith, Keemia Azvine, Maite Arribas, Jiaqi Zhou, Serena Deiana, Bastian Hengerer, Emma N Cahill

## Abstract

Fear and anxiety are adaptive states that allow humans and animals alike to respond appropriately to threatening cues in their environment. Commonly used tasks for studying behaviour akin to fear and anxiety in rodent models are pavlovian threat conditioning and the elevated plus maze (EPM) respectively. In threat conditioning the rodents learn to associate an aversive event with a specific stimulus or context. The learnt association between the two stimuli (the ‘memory’) can then be recalled by re-exposing the subject to the conditioned stimulus. The elevated plus maze is argued to measure the agoraphobic avoidance of the brightly lit open maze arms in crepuscular rodents. These two tasks have been used extensively, yet research into whether they interact is scarce. We investigated whether recall of an aversive memory, across contextual, odour or auditory modalities, would potentiate anxiety-like behaviour in the elevated plus maze. The data did not support that memory recall, even over a series of timepoints, could influence EPM behaviour. Furthermore, there was no correlation between EPM behaviour and conditioned freezing in independent cohorts tested in the EPM before or after auditory threat conditioning. Further analysis found the production of 22kHz ultrasonic vocalisations revealed the strongest responders to a conditioned threat cue. These results are of particular importance for consideration when using the EPM and threat conditioning to identify individual differences, and the possibility to use the tasks in batteries of tests without cross-task interference.

## INTRODUCTION

Tasks to measure rodent behaviour motivated by aversive experience are commonly used to model features of symptoms present in anxiety disorders. Nonetheless, there is much discussion on how to best describe and validate what the tasks are detecting in terms of emotionally driven behaviour, and what the tasks might predict for translation (for review see Bach, 2021; Steimer, 2011). An influential system of classification of rodent defensive responses places the response along a theoretical axis of psychological or physical predator imminence versus escapability of the context (Fanselow et al., 1988). The predator imminence model has been used to categorise specific readouts, such as freezing or avoidance, as more akin to states of fear or anxiety respectively (McNaughton and Corr, 2004, Blanchard and Blanchard, 1969). However, whether a clear distinction between emotional states of fear or anxiety is fully warranted for effective translation has recently come under debate given the overlap in mechanisms recruited by behaviour related to each (Daniel-Watanabe and Fletcher, 2021).

In rodents, innate avoidance of open arm exploration in the elevated plus maze (EPM) has been argued to reflect an anxiety-like agoraphobia (Walf and Frye, 2007). The production of 22kHz ultrasonic vocalisations, or so-called alarm calls, has been posited to also reflect a state akin to anxiety (Anderson, 1954; Schwarting and Wöhr, 2012). Interestingly, the production of 22kHz is not typically reported in the EPM (Borta et al., 2006), but is more reliably triggered by exposure of rodents to a conditioned threat stimulus (CS) or mild shocks such as during pavlovian conditioning (Wöhr and Schwarting, 2008). Conditioned freezing to threat cues has more recently been rebranded from a description as a fear-related behaviour (LeDoux, 2014), but is often classed as distinct from the anxiety-like behaviours recorded in maze tasks such as the EPM. Despite widespread use of the EPM and conditioned freezing tasks, sometimes in tandem, there is little consensus on whether the tasks do measure truly independent behaviours or indeed how recent experience impacts performance in these tasks (File, 1993). One could hypothesize that recall of a recent aversive experience would lead to heightened anxiety responses within the window of that memory reactivation. This hypothesis is consistent with the notion that the elevated plus-maze is sensitive to acute behavioural states.

On the other hand, the EPM has been used as a measure of so-called trait-like, anxiety, which suggests it could be stable over time and potentially reveal individual differences that translate into resilience or risk for pathological responses (Shumake et al., 2014; Richter-Levin et al., 2019). Considering the rising debate on the distinction of rodent defence responses as fear-like or anxiety-like, we examined whether a potential interaction between unconditioned behaviour in the EPM and conditioned freezing may exist. In addition, we report on the production of 22kHz USVs in response to auditory threat conditioning cues. To better characterize potential impacts of prior aversive experience on EPM performance a series of experiments were carried out that systematically varied the time since stimulus presentation and the modality of the conditioning stimulus. Furthermore, we examined whether the EPM measurements taken days before or after threat conditioning measurements held any relationship.

## MATERIALS AND METHODS

### Subjects

Subjects were 104 adult male Lister-Hooded rats (Charles River and Envigo, UK) weighing approx. 250–300g at the start of experiments. All animals were housed in groups of four per cage and kept under a reversed 12 h light/dark cycle (lights off 07:00 h until 19:00 h) and were provided with food and water ad libitum, except for during behavioral procedures, which were conducted during the rats’ dark cycle. The rats were randomly assigned to each group and experimenters were blind to group allocation for subsequent analysis. This research was conducted on Project Licence 70/7548 and has been regulated under the Animals (Scientific Procedures) Act 1986 Amendment Regulations 2012 following ethical review by the University of Cambridge Animal Welfare and Ethical Review Body (AWERB).

### Behavioral procedures

For conditioned freezing, four identical operant boxes (29.5 × 32.5 × 23.5cm, MedAssociates) were used with a Plexiglass rear wall, hinged door and roof, with aluminium side walls. The boxes also contained a house light (2.5W, 24V), a speaker (3kHz tone, 75 dB), ultrasonic microphones (Metris, Netherlands) and a CCTV video camera (Spy Camera model CMOSNC76). The boxes were positioned within sound attenuating chambers and each also contained a ventilation fan, which provided background noise (approx. 60dB). At time zero the house light was turned on. The same box was used for each rat throughout the experiments. The unconditioned stimulus was a mild foot-shock (0.5mA, 0.5s). All training and test sessions were video recorded for off-line behavioural analysis. The percentage of time freezing (absence of movement except for breathing) during 1 min before (Pre CS) and during the CS was manually scored from the videos by observers blind to the groups.

### Contextual Conditioning

Rats were individually placed in the conditioning boxes, which they had not previously been habituated to. A mild electric foot shock (0.5mA, 0.5s) was administered after the rats had been in the context for 2 min to allow the contextual representation to first be encoded (Fanselow, 1990). The context-shock pairing was repeated 2 more times with 2 min intervals between. Following the last shock and 2 min interval, the house light remained on for a further 2 min. Following this, the house light was turned off and the rats were removed from the conditioning boxes and then returned to their home cage. No Shock controls were placed in the context for the same duration. For the recall test, rats were placed in the context for 3 mins.

### Olfactory Conditioning

Rats were individually habituated to the context for 1 hour on Day 0 and Day 1. Rats were then conditioned to associate the foot shocks with the odour acetophenone (Sigma Aldrich). The odour, acetophenone, was prepared in a separate room and gloves were changed by the experimenters before handling the rats. The odour was diluted to 10% in mineral oil and 200µl of the diluted odour was placed onto a cotton pad. Following a 4-minute baseline period, the odour was introduced into the shock box in the waste tray and the conditioned group received three foot shocks with a 2 min inter-trial interval. The control odour exposure group (No Shock) was exposed to the odour for the same duration but did not receive foot shocks. Between sessions a fan was run for 15 mins to ventilate the room. For the recall test, rats were placed back in the context and after three mins had passed the odour was introduced on a cotton pad as described above. The rats remained in presence of the odour for 3 mins.

### Auditory Conditioning

On Day 0 rats were habituated to the context for 1 hour. On Day 1, after a baseline period of 20 min three pairings of the 1 min tone (3kHz, 75dB) coterminous with the footshock were presented with 5 min inter-stimulus intervals. At recall test, after 5 min baseline the tone was presented for 1 min and then the rats remained in the context for a further 5 mins.

### Elevated Plus-maze

Each rat was tested individually on a plus-maze made from black Perspex with four arms of 50 cm long and 10 cm wide, at a height of 50 cm from the floor, with raised sides of 30 cm (ViewPoint, France). The maze was situated in a room with many external cues located around the maze, and these cues remained the same throughout training and testing of each cohort. At the start of each session, a rat was placed on the end of a same open arm facing away from the middle region. The experimenter triggered the recording and left the room to observe from an adjacent room. After 5 min the video recording was stopped, and the experimenter re-entered the room and removed the rat from the maze. Rats were then returned to their home cage. Between sessions, the maze was cleaned with water and dried. In the singular case where a rat jumped off the maze, the experimenter re-entered the room, returned the rat to the home cage and this rat was excluded from statistical analyses. Time spent in the open and closed arms and in the centre zone was manually scored and a percentage of the total time was calculated. In addition, the absolute number of entries into open and closed arms and the latency to first enter the open arms were scored. The rat was scored to have entered a particular arm when its hind legs passed the border.

### USV Analysis

An Ultrasound Microphone (Metris, Netherlands) was placed through a hole in the middle of the operant box roof about 30 cm above the shock floor. The microphones were sensitive to frequencies of 15–125 kHz. Vocalization was recorded and analysed using the Sonotrack software (Metris, Netherlands). Call detection was provided by an automatic threshold-based algorithm. Experimenters manually screened the calls detected and classified based on the mean frequency as 22kHz calls or 55+kHz calls or cage noise. The number of USV calls and total calling time were analysed for the 22kHz calls only.

### Statistical Analysis

Data are presented as mean ± SEM, unless otherwise stated. Statistical analyses included t-tests, repeated-measures ANOVA, and planned Sidak comparisons for more than three groups were made using GraphPad Prism (GraphPad Software Inc., La Jolla, CA, USA, version 9.3.0). Where Mauchly’s Test of Sphericity indicated the assumption of sphericity had been violated, degrees of freedom were Greenhouse-Geisser corrected. Where there was a clear prediction for the direction of an effect a one-tailed t-test was performed. For comparison of categorical data, i.e. the proportion of rats that produced 22kHZ vocalisation or not, a Fisher’s Exact Test was performed. The significance level was set at p<0.05. Graphs and figures were generated in GraphPad Prism 9.3.0.

## RESULTS

### Recall of a pavlovian aversive memory had no impact on subsequent elevated plus-maze behaviour

#### Contextual Conditioning

Whether recall of an aversive experience would impact behaviour in the EPM tested shortly after memory recall was assessed. The responses of a contextual threat conditioned group (Shock) were compared to a control group of rats that were exposed to the context for the same duration of time but did not receive any foot shocks (No Shock, Fig 1). As expected, at the context recall test the shocked group showed a strong conditioned freezing response relative to the No Shock controls (one-tailed t-test, t_10_=4.807, P=0.0004). 90 minutes later, all rats were tested for arm exploration and entries in the EPM. The time spent in the closed arms was not different between the groups (one-tailed t-test, t_10_=1.024, P=0.1651). Although one rat in the Shock group did not enter the closed arm after initiation of the test at all, and showed some intermittent freezing responses in the open arm, we did not exclude the data as a statistical outlier as exclusion did not influence the conclusion of the findings. Previous results in the literature have demonstrated that 90’ after stress or recall of a stressful experience EPM behaviour was negatively impacted (Mechiel Korte et al., 1999). Using independent conditioned groups, a series of time points after recall were further examined to see if any temporal impact of memory recall on the EPM could be detected (Fig 2). None of the time points examined influenced the time spent in the closed arms, most notably even for rats that were introduced to the EPM as directly as possible after context recall (0’ group).

**Figure 1:**
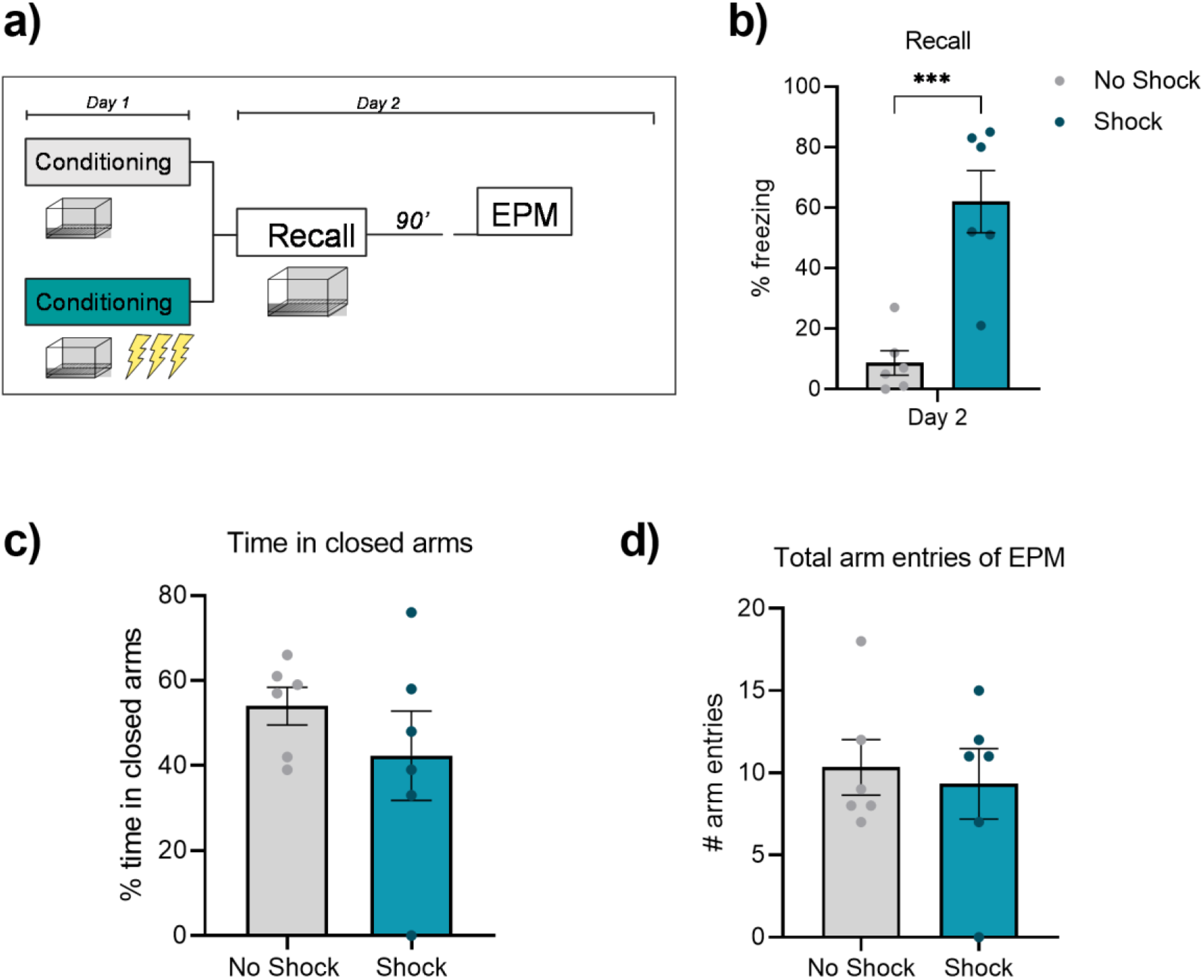
Recall of contextual threat memory does not influence subsequent EPM behaviour. a) Rats were either threat conditioned by pairing mild footshocks with a contextual environment (Shock), or placed in the context in absence of footshock (No Shock). A memory recall test was performed and then 90’ later they were tested in the EPM. b) Only the Shock group showed conditioned freezing at recall of the context. c) 90’ later, the time spent in the closed arms of the EPM was equivalent between the two groups. D) Total arm entries were equivalent between the groups.

**Figure 2:**
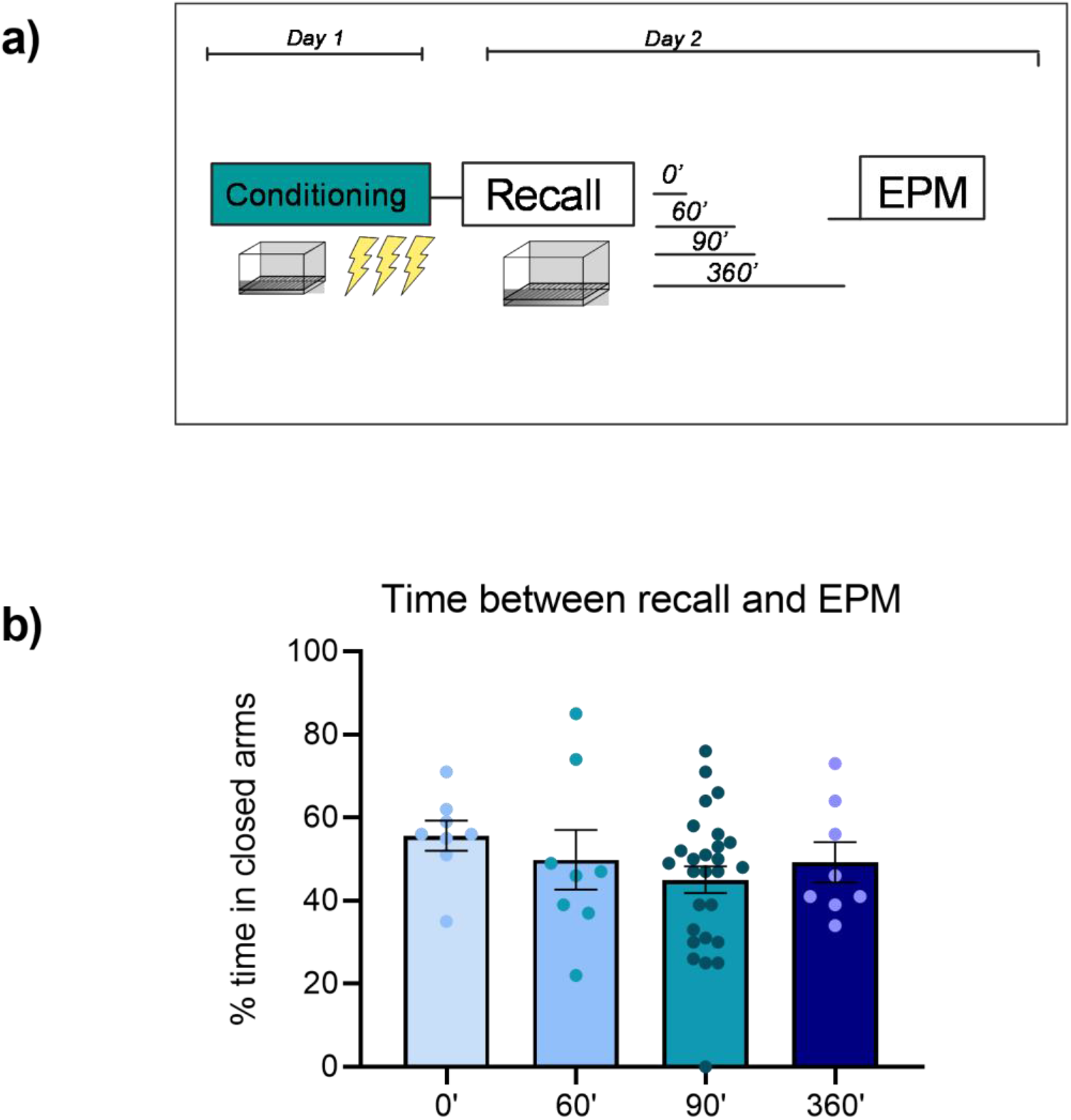
Temporal analysis of aversive context recall on EPM behaviour. a) Independent cohorts were tested at a series of time points after recall on conditioned threat context memory to investigate any temporal effect of threat memory recall on EPM measures. b) None of the timepoints investigated showed increased closed arm exploration after threat memory recall.

#### Olfactory Conditioning

It could be argued that a contextual conditioning protocol might be prone to extinction during the recall test of the memory, as the cue (ie. the context) is consistently present during the recall trial in absence of the US. Although no significant within session extinction during contextual recall was observed, we proceeded to test if a more discrete modality of aversive memory recall would impact the EPM behaviour. Pavlovian olfactory conditioning was performed and a control group (No Shock) were exposed to the neutral odour for the same duration as those that received paired-footshocks (Shock, Fig 3). To check that exposure to the odour during recall would not result in complete extinction of the association, a second recall test was performed eight days later and the rats were tested in the EPM 90’ after that recall session. Significant levels of freezing were detected in the Shock group, relative to the No Shock group (F_1,10_= 62.24, P<0.001) - and to pre-CS contextual levels of freezing (data not shown)-still at the Day 9 second recall test (P= 0.0169). Both groups were tested in the EPM 90 min after recall. There was no significant difference between the time spent in the closed arms of the EPM nor total arm entries. This suggests that recent recall of an olfactory aversive memory does not impact subsequent behaviour in the elevated plus-maze.

**Figure 3:**
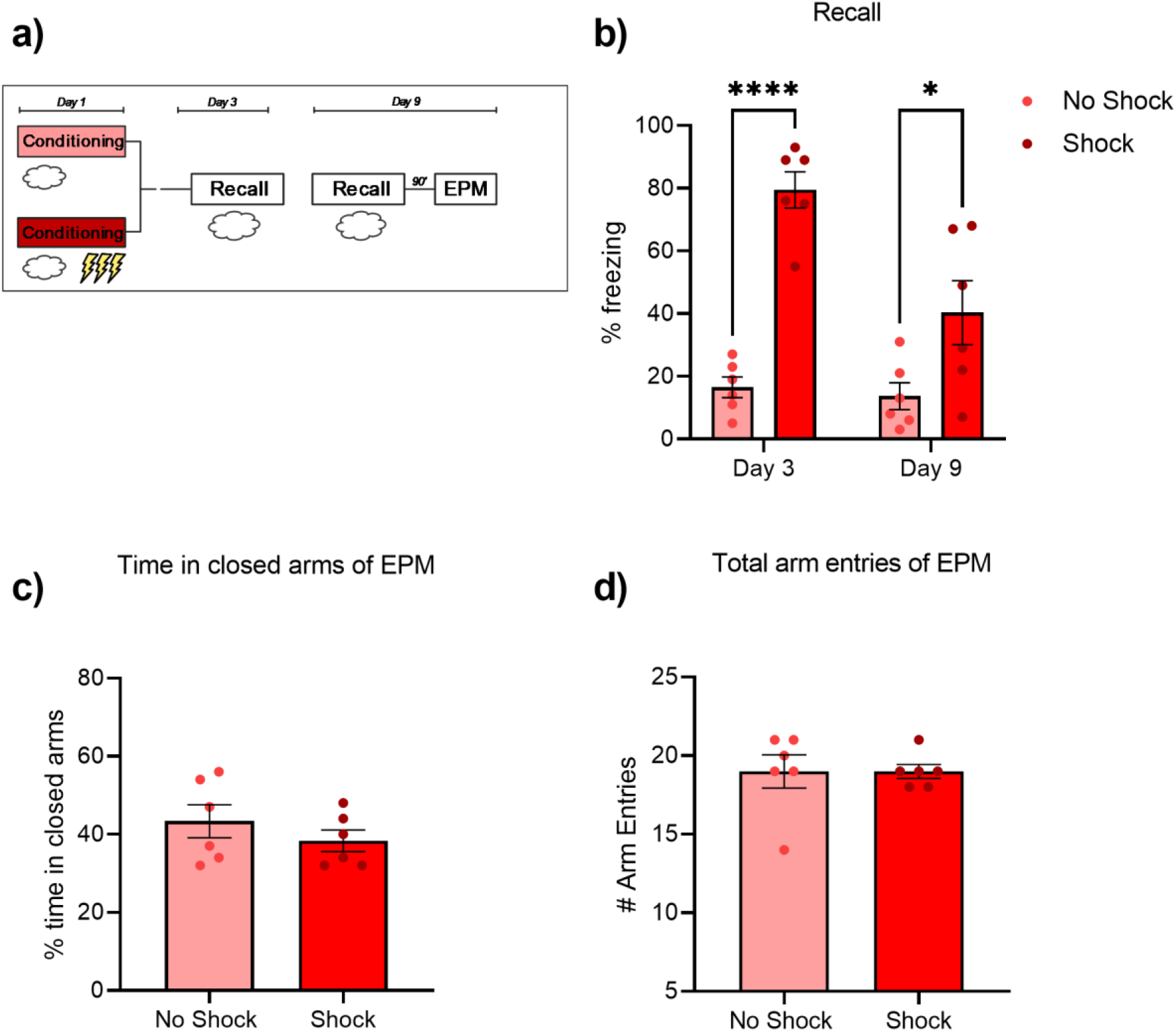
Recall of olfactory CS memory does not influence EPM behaviour 90 mins later. a) Rats were conditioned by pairing mild footshocks with a neutral odour acetophenone (Shock), or presented the odour in absence of footshock (No Shock). b) An initial test of consolidation was performed at Day 3, followed by a subsequent test at Day 9 to assess the stability of the olfactory cue memory. The Shock group showed persistent conditioned freezing at recall to the odour CS. c) 90’ after memory recall the time spent in the closed arms of the EPM was equivalent between the two groups. D) Total arm entries were equivalent between the groups.

#### Auditory Conditioning

The final modality of conditioning examined was to use an auditory tone as the CS. As before, a control group was presented the tone in absence of footshocks (No Shock). A conditioned group (Shock) were trained with paired tone footshock presentations (Fig. 4). The next day a recall test was performed 90’ before testing in the EPM. As seen with the contextual and olfactory modalities, the conditioned freezing was specific to the paired presentation (t_19_ =5.009, P<0.0001). The tone memory recall had no impact on any of the EPM measures.

**Figure 4:**
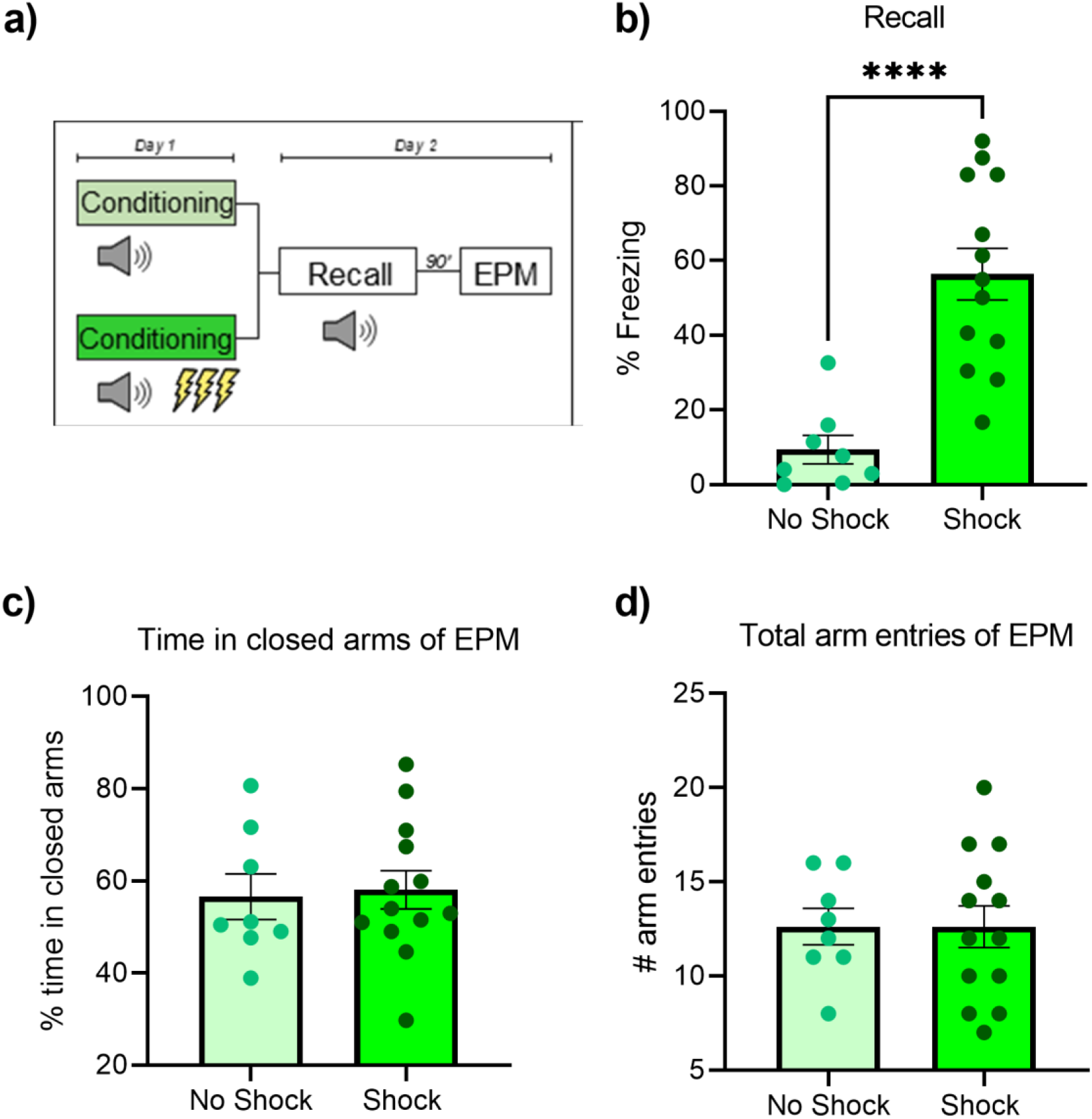
Recall of tone CS memory does not influence EPM behaviour 90 mins later. a) Rats were conditioned by pairing mild footshocks (Shock) with an auditory tone, or presented the tone in absence of footshock (No Shock). A memory recall test was performed and then 90’ later they were tested in the EPM. b) The Shock group showed conditioned freezing at recall to the CS. C) Time spent in the closed arms of the EPM was equivalent between the two groups. D) Total arm entries were equivalent between the groups.

The independence of EPM measures from experience of conditioned threat was further tested across a longer time period. We examined whether the time spent in the closed arm related to CS-evoked freezing, or *vice versa* whether CS-evoked freezing would relate to subsequent EPM behaviour in two independent cohorts (Fig. 5). In one cohort, rats were screened in the EPM to examine if this behaviour related to subsequent conditioned freezing. At the recall test of auditory cued-conditioning, eight days after the EPM test, there was no relationship between time spent in the closed arm and the freezing during the test CS presentation (r(9)=0.2934, p=0.4107). In another cohort, the order of testing was reversed, and rats underwent auditory cue-conditioning then seven days later they were tested in the EPM. Conditioned freezing at test during the CS presentation did not relate to subsequent closed arm time in the EPM (r(9)= -0.1546, p=0.6698). There were no significant differences in the mean levels of CS freezing nor time in the closed arms between the cohorts.

**Figure 5:**
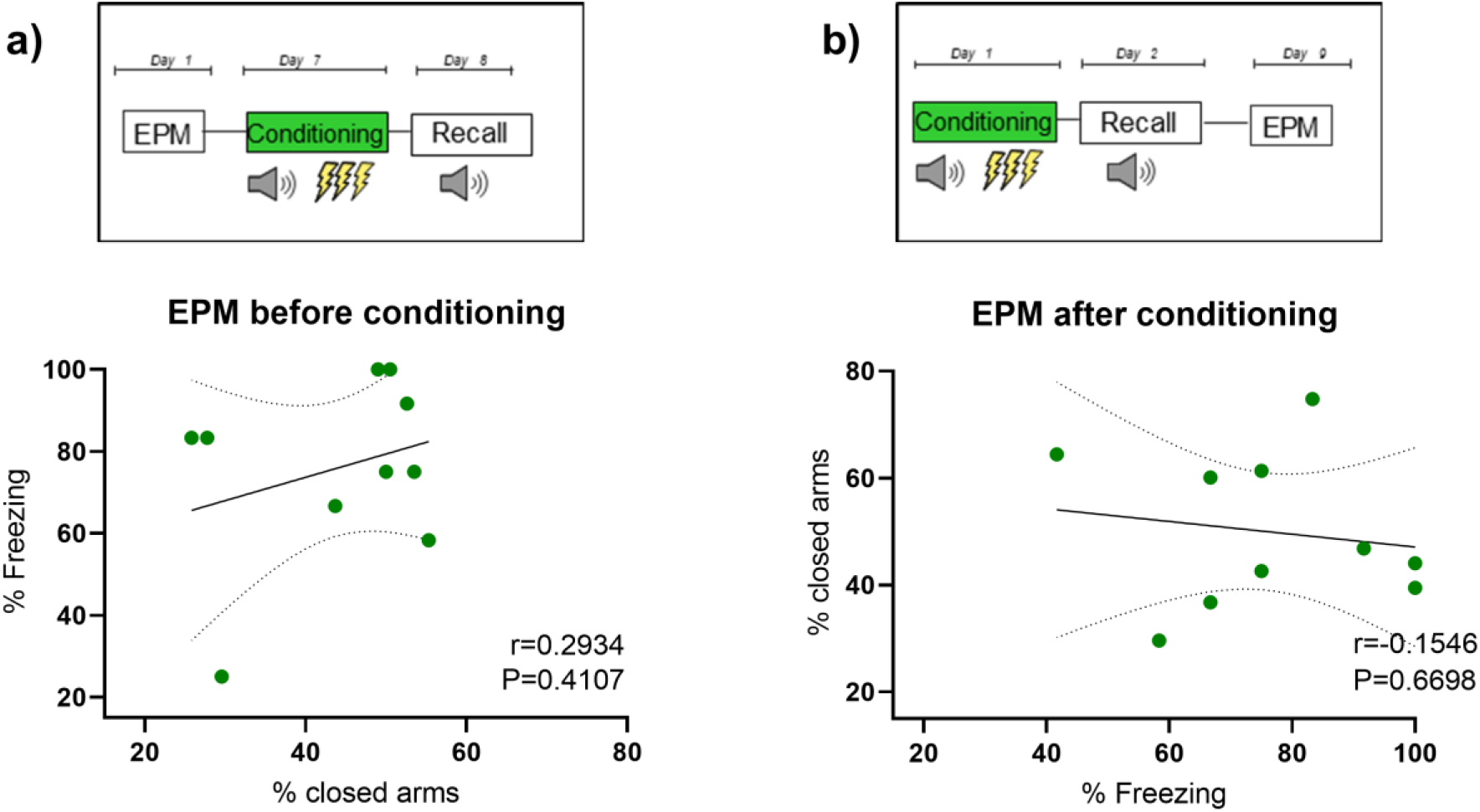
Time spent in the closed arm of the EPM does not relate to CS-test conditioned freezing. a) The time spent in the closed arm of the EPM did not relate to subsequent conditioned freezing at tone CS-test 8 days later. b) Conditioned freezing at CS-test did not relate to the time spent in the closed arm of the EPM 7 days later. N=10 for each independent cohort.

**Figure 6:**
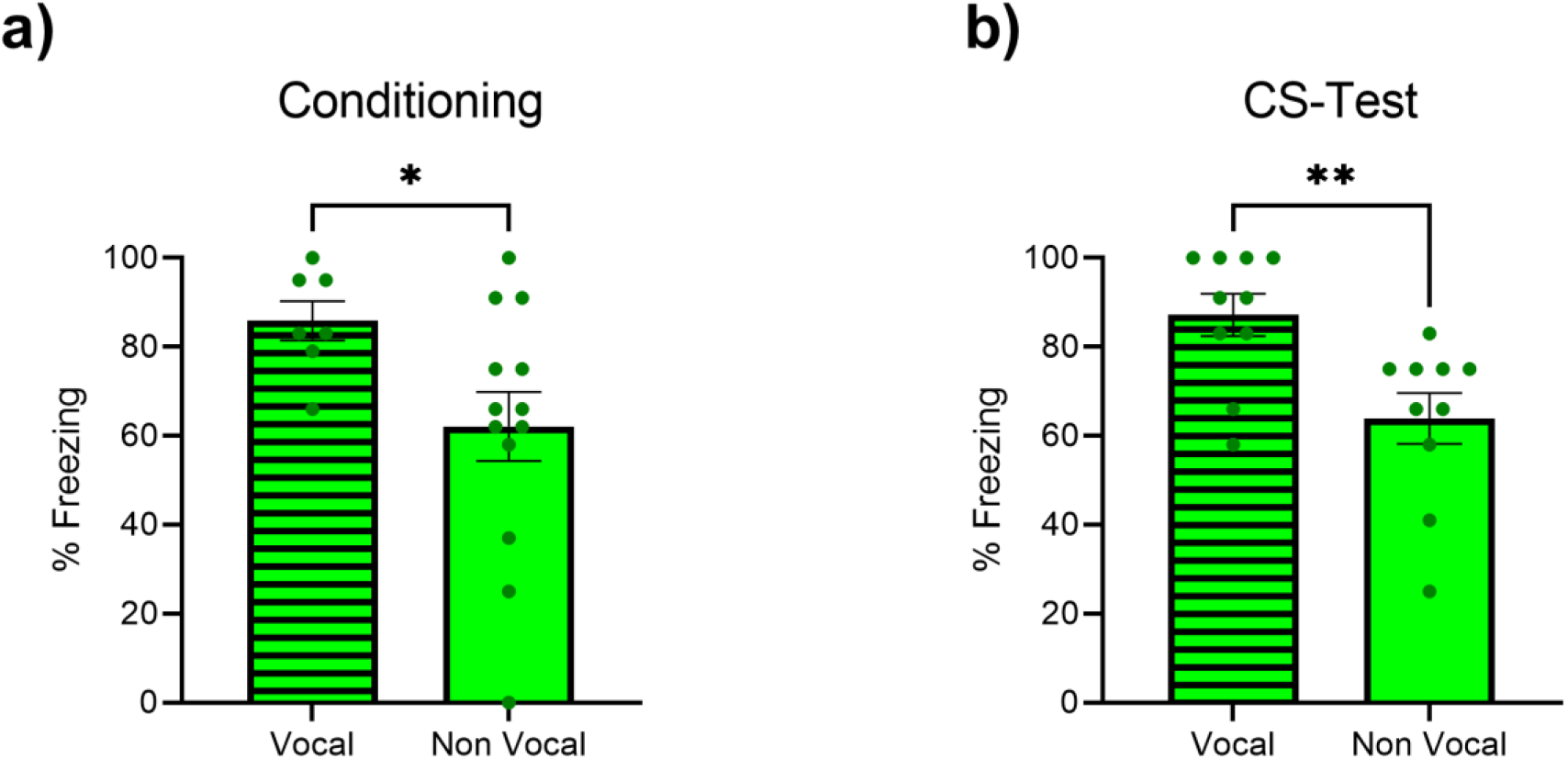
22kHz vocalisation reveals the strongest responders during conditioning and recall. a) Rats that produced 22kHz vocalisations had higher levels of freezing by the end of conditioning than those not producing calls. b) 24h later, the rats that produced 22kHz vocalisations in response to the tone CS at recall were those with significantly higher levels of freezing.

Taken together these findings suggested that the behaviour in the EPM could be read as an innate behaviour measure independent of recent aversive experience, as it did not relate to conditioned threat detection behaviour (i.e. conditioned freezing) under any of the tested modalities nor timepoints.

#### Rats that produced 22kHz USVs displayed stronger conditioned freezing

The production of 22kHz USVs calls during the auditory threat cue conditioning and CS-test sessions were analysed in both cohorts of rats screened with the EPM before or after cue conditioning. In the cohort tested in the EPM before cue conditioning, the time spent in the closed arm did not correlate with the total duration of USV calls through subsequent conditioning nor at CS-test. Also, the durations of USVs made during conditioning or during the CS-test of the cohort examined with the EPM afterwards did not relate to closed arm behaviour at that time. Whilst it must be acknowledged that there is considerable inter-individual variability in the propensity to produce USVs, from the present data the performance in the EPM did not seem to reflect the presence or absence of calling during conditioned freezing. On the other hand, we observed that the production of 22kHz USVs was tightly coupled to conditioned freezing behaviour and so hypothesized these rats would show the strongest freezing response to the threat cue. In both cohorts, rats that produced USVs displayed higher levels of freezing than those that did not vocalise during CS-conditioning (one tailed t-test, T_18_=2.127, P=0.0238). At CS-test, the difference in freezing between vocal and non-vocal rats was also significant (one tailed t-test, T_18_=3.138, P=0.0028). As such the production of 22kHz calls triggered by a threat cue reveals the heightened state of the most reactive individuals (Fig. 2).

## DISCUSSION

Presentation of an aversive conditioned stimulus (CS) at a recall test has been shown to activate brain circuits shared with those activated by EPM experience, somewhat depending on the modality of the associative stimulus (Steimer, 2002; Tovote et al., 2015). Therefore, it is possible that there may be some overlap between the brain mechanisms recruited by experience of the EPM and threat conditioning (Wang et al., 2011; Zhang et al., 2014; Jimenez et al., 2018). Although the conditioning modalities examined herein, namely contextual, olfactory and auditory, recruit stimulus-specific circuits they have all been shown to require intact amygdala function for the associative learning of unconditioned stimulus to CS pairings (Kim and Fanselow, 1992; Cousens and Otto, 1998). Thus, the potential interaction between the mechanisms underlying EPM behaviour and conditioned freezing makes it interesting to compare how these two measures may relate to each other.

The series of experiments described herein established that the EPM was insensitive to recent aversive memory recall and the exploratory behaviour seen in the EPM did not relate to conditioned freezing, across the modalities examined. These findings might be considered surprising given that others have reported negative effects on open arm activity, sometimes taken as a measure of ‘state anxiety’, following re-exposure to contexts associated with footshocks (Mechiel Korte et al., 1999; Marinzalda et al., 2014). Although Mechiel Korte and colleagues did not report conditioned freezing data, it is unlikely that the lack of ‘fear-potentiation’ effect of context re-exposure reported here was simply due to some weaker associative learning because no correlation was detected between freezing at memory recall and the time spent in the closed arms across the experiments, where a range of strength of conditioned responding was seen at the recall test. Most notably the levels of freezing induced by olfactory conditioning were stronger at the first recall test than those elicited by auditory or contextual cues paired with equivalent level and number of footshocks. It could be argued that the aversive state induced by recall of aversive memory is perhaps too weak to impact EPM behaviour, yet others have reported similar observations even with a relatively strong (controllable or uncontrollable) tailshock, in that they did not influence EPM arm exploration measures 2 hours later (Grahn et al., 1995). Of course it is possible that methodological differences in EPM conditions may underlie the different findings (Wall and Messier, 2001), nonetheless the levels of exploration seen herein do not differ vastly from those reported elsewhere in the literature. A possible interpretation of these findings is then that the EPM is not acutely sensitive to recall of an aversive experience and measures a more trait-like level of open arm exploration.

Given an interpretation that the EPM measures may reflect something more akin to a stable trait measure, two cohorts were screened in the EPM prior to or after conditioning to see if closed arm exploration would correlate to the acquisition or expression of conditioned freezing or *vice versa* across a longer time window. Differences in conditioned freezing recall test have been reported to be revealed by performing median splits of rats into the extreme of responses on open arm (OA) activity performed around a month before the recall test (Borta et al., 2006). In agreement with the prior study (Borta et al., 2006), when we performed a median split of the group EPM tested a week before threat conditioning the low OA (LOA) explorative rats had significantly higher levels of freezing behaviour during the CS presentation of the recall test session (one tailed t-test, T_8_ = 0.288 P=0.0351) despite equivalent freezing during conditioning training. However, when taken as a whole group the arm exploration values did not significantly correlate to freezing behaviour. We also performed the same split analyses of the cohort tested with the EPM one week after cued-threat conditioning. The levels of freezing during conditioning or during the CS-test did not differ between the subsequently segregated LOA and High OA groups (P=0.1788 and P=0.3495 respectively). Taken together, whilst there seemed to be some promise in a median split of OA behaviour before threat conditioning to relate to the conditioned freezing at test, the cohort split by OA behaviour after threat conditioning indicated no differences in their prior conditioned freezing. Due to the narrow propensity of the split of EPM behaviour to relate to subsequent conditioned freezing, our analyses were performed without any median split segregation of the rats by EPM behaviour to capture the full range of responses. Studies have also demonstrated that rats that were poor at discrimination of a CS+ from a CS-at a recall test were statistically prone to spend more time in the closed arms of the 2 days after the recall (Duvarci et al., 2009). These findings might suggest together that in the extremes of memory precision the EPM may reveal meaningful individual differences. Overall, there appears to be limited relationship of the EPM measures to conditioned freezing in the full range of responses taken without thresholds to segregate groups of rats based on responses.

Throughout the experiments the production of 22kHz USVs was consistently seen to be present in the rats showing the strongest conditioned freezing responses. These two behaviours, in contrast to arm avoidance in the EPM and freezing, seem to be closely tied together. The interplay of 22kHz USVs and conditioned freezing represents an interesting means to reveal the most reactive individuals in a threat conditioning task. If freezing is taken as an active ‘fear-like’ response to evade detection by an inescapable predator (McNaughton and Corr, 2004), the progression to 22kHz calls by an individual in isolation may represent a shift to an active response to deter a predator stalking attack (Blanchard et al., 1991). Notably, the levels of USVs did not correlate to prior or subsequent EPM behaviour, again tying the production of such alarm calls to sensitive conditioned threat cue detection. Nonetheless, the variability in production of USVs again points to important individual differences in propensity to react to a threat cue.

Variations of mazes that expose rodents to open areas are routinely used to measure behaviour argued to be akin to anxiety symptoms (Cryan and Holmes, 2005). Interestingly in mice there is evidence that whilst the open field task correlated to levels of conditioned contextual freezing, the EPM did not correlate to either baseline nor conditioned contextual freezing (Ahn et al., 2013), which mirrors our findings here with auditory conditioning. In the same study it was also remarked that the performance in the open field task did not correlate with the EPM behaviour, perhaps surprising given how these tasks are thought to both track an agoraphobia driven behaviour, but other work has substantiated these differences in measures taken in each task (Carola et al., 2002). Indeed, in the early studies of ‘fear-like’ responses that preceded the now widespread use of a traditional EPM set-up, Montgomery argued in a series of papers a distinction between fear and exploratory behaviour in mazes that was driven more by novelty than ‘fear’ (Montgomery and Monkman, 1955). Despite the potential issues with the EPM as a translational task (Ennaceur, 2014), the independence of measures in the EPM from conditioned freezing may simply indicate that they measure different forms of aversively motivated behaviours that are context appropriate. This may be advantageous for studies that use the tests in series in a battery of tasks. Given the prospect that individual differences in performance at the extremes proves interesting for further exploration, these presented findings call for caution in generalisation of interpretations from the typical range of behaviour in the EPM across other aversively motivated tasks.

## FUNDING ACKNOWLEDGMENTS AND DISCLOSURE

This work was funded by a collaborative research grant with Boehringer Ingelheim Pharma GmbH & Co. awarded to E.N.C.

## REFERENCES

Ahn SH, Jang EH, Choi JH, Lee HR, Bakes J, Kong YY, Kaang BK (2013) Basal anxiety during an open field test is correlated with individual differences in contextually conditioned fear in mice. Animal Cells Syst (Seoul) 17:154–159.

Anderson JW (1954) The Production of Ultrasonic Sounds by Laboratory Rats and Other Mammals. Science 119:808–809.

Bach DR (2021) Cross-species anxiety tests in psychiatry: pitfalls and promises. Mol Psychiatry Available at: http://dx.doi.org/10.1038/s41380-021-01299-4.

Blanchard RJ, Blanchard DC (1969) Crouching as an index of fear. Journal of Comparative and Physiological Psychology 67:370–375.

Blanchard RJ, Blanchard DC, Agullana R, Weiss SM (1991) Twenty-two kHz alarm cries to presentation of a predator, by laboratory rats living in visible burrow systems. Physiol Behav 50:967–972.

Borta A, Wöhr M, Schwarting RKW (2006) Rat ultrasonic vocalization in aversively motivated situations and the role of individual differences in anxiety-related behavior. Behav Brain Res 166:271–280.

Carola V, D’Olimpio F, Brunamonti E, Mangia F, Renzi P (2002) Evaluation of the elevated plus-maze and open-field tests for the assessment of anxiety-related behaviour in inbred mice. Behav Brain Res 134:49–57.

Cousens G, Otto T (1998) Both pre-and posttraining excitotoxic lesions of the basolateral amygdala abolish the expression of olfactory and contextual fear conditioning. Behav Neurosci 112:1092–1103.

Cryan JF, Holmes A (2005) Model organisms: The ascent of mouse: Advances in modelling human depression and anxiety. Nat Rev Drug Discov 4:775–790.

Daniel-Watanabe L, Fletcher PC (2021) Are Fear and Anxiety Truly Distinct? Biol Psychiatry Glob Open Sci:1–9 Available at: https://doi.org/10.1016/j.bpsgos.2021.09.006.

Duvarci S, Bauer EP, Paré D (2009) The bed nucleus of the stria terminalis mediates inter-individual variations in anxiety and fear. J Neurosci 29:10357–10361.

Ennaceur A (2014) Tests of unconditioned anxiety - Pitfalls and disappointments. Physiol Behav 135:55–71

Fanselow M, Angeles L, Lester LS (1988) A Functional Behavioristic Approach to Aversively Motivated Behavior: Predatory Imminence as a Determinantof the Topography of Defensive Behavior. Evol Learn:197–224.

File SE (1993) The interplay of learning and anxiety in the elevated plus-maze. Behavioural Brain Research 58:199–202

Grahn RE, Kalman BA, Brennan FX, Watkins LR, Maier SF, Grahn RE, Kalman BA, Brennan FX, Watkins LR, Maier SF (1995) The Elevated Plus-Maze Is Not Sensitive to the Effect of Stressor Controllability in Rats. Pharmacology, Biochemisty and Behaviour 52:565–570

Jimenez JC, Su K, Goldberg AR, Luna VM, Biane JS, Ordek G, Zhou P, Ong SK, Wright MA, Zweifel L, Paninski L, Hen R, Kheirbek MA (2018) Anxiety Cells in a Hippocampal-Hypothalamic Circuit. Neuron 97:670–683.e6.

Kim JJ, Fanselow MS (1992) Modality-specific retrograde amnesia of fear. Science 256:675–677.

LeDoux JE (2014) Coming to terms with fear. Proc Natl Acad Sci U S A 111:2871–2878

Marinzalda MDLA, Pérez PA, Gargiulo PA, Casarsa BS, Bregonzio C, Baiardi G (2014) Fear-potentiated behaviour is modulated by central amygdala angiotensin II A T1 receptors stimulation. Biomed Res Int 2014:1–7.

McNaughton N, Corr PJ (2004) A two-dimensional neuropsychology of defense: Fear/anxiety and defensive distance. Neurosci Biobehav Rev 28:285–305.

Mechiel Korte S, De Boer SF, Bohus B (1999) Fear-potentiation in the elevated plus-maze test depends on stressor controllability and fear conditioning. Stress 3:27–40.

Montgomery KC, Monkman JA (1955) The relation between fear and exploratory behavior. J Comp Physiol Psychol 48:132–136.

Richter-Levin G, Stork O, Schmidt M V. (2019) Animal models of PTSD: a challenge to be met. Mol Psychiatry 24:1135–1156

Schwarting RKW, Wöhr M (2012) On the relationships between ultrasonic calling and anxiety-related behavior in rats. Brazilian J Med Biol Res 45:337–348.

Shumake J, Furgeson-Moreira S, Monfils MH (2014) Predictability and heritability of individual differences in fear learning. Anim Cogn 17:1207–1221.

Steimer T (2002) The biology of fear-and anxiety-related behaviors. Dialogues Clin Neurosci 4:231–249.

Steimer T (2011) Animal models of anxiety disorders in rats and mice: some conceptual issues. Dialogues in Clin Neurosci 13:495–506

Tovote P, Fadok JP, Lüthi A (2015) Neuronal circuits for fear and anxiety. Nat Rev Neurosci 16:317–331.

Walf AA, Frye CA (2007) The use of the elevated plus maze as an assay of anxiety-related behavior in rodents. Nat Protoc 2:322–328.

Wall PM, Messier C (2001) Methodological and conceptual issues in the use of the elevated plus-maze as a psychological measurement instrument of animal anxiety-like behavior. Neurosci Biobehav Rev 25:275–286.

Wang D V., Wang F, Liu J, Zhang L, Wang Z, Lin L (2011) Neurons in the amygdala with response-selectivity for anxiety in two ethologically based tests. PLoS One 6:1–7.

Wöhr M, Schwarting RKW (2008) Ultrasonic calling during fear conditioning in the rat: no evidence for an audience effect. Anim Behav 76:749–760.

Zhang WN, Bast T, Xu Y, Feldon J (2014) Temporary inhibition of dorsal or ventral hippocampus by muscimol: Distinct effects on measures of innate anxiety on the elevated plus maze, but similar disruption of contextual fear conditioning. Behav Brain Res 262:47–56.

